# Too little, too late: transcription during imbibition of lethally aged soybean seeds is weak and delayed, but not aberrant

**DOI:** 10.1101/2021.03.25.437023

**Authors:** Margaret B. Fleming, Eric L. Patterson, Christina Walters

**Affiliations:** USDA-ARS, National Laboratory for Genetic Resource Preservation, 1111 S. Mason Street, Fort Collins, CO, 80521, USA; Department of Plant Biology, 612 Wilson Road, Room 262, Michigan State University, East Lansing, MI 48824; Department of Plant, Soil and Microbial Sciences, Giltner Hall, 293 Farm Lane, Room 103, East Lansing, MI 48824

**Keywords:** aging, gene expression, germination, imbibition, metabolism, mortality, seed storage, viability

## Abstract

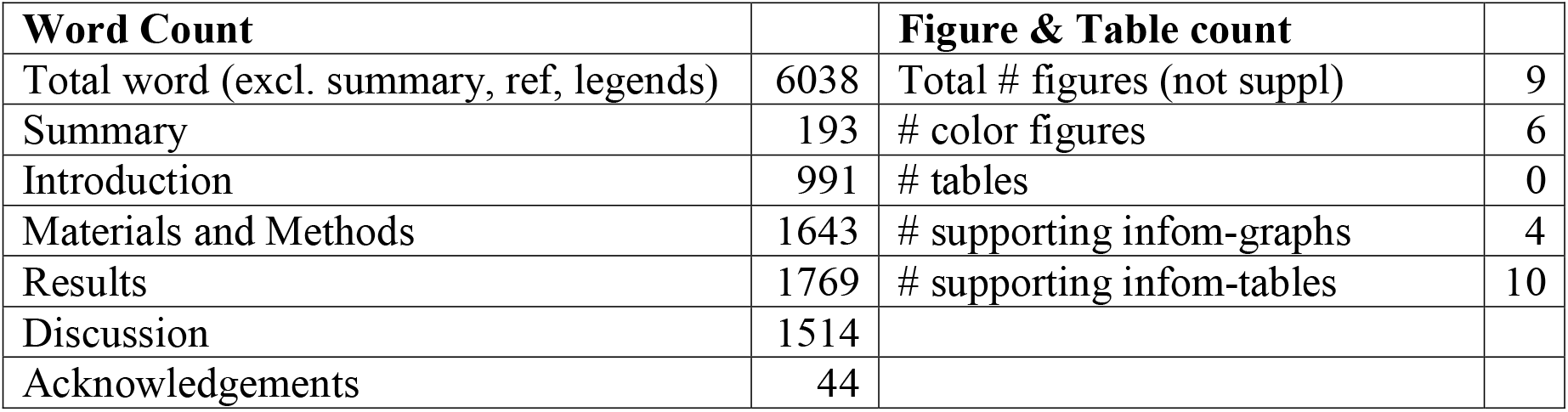

- This study investigates alive to dead signals in seeds that aged during cool, dry storage. Signals may invoke abrupt, lethal metabolic pathways or reflect effects of accumulated small injuries which impair recovery from life in the dry state.
- Cohorts of soybean (*Glycine max* cv. ‘Williams ’82) seeds were stored for 3, 19 and 22 years. Transcriptomes of dry embryonic axes and axes 24 hours after imbibition (HAI) were sequenced to determine gene expression patterns. These cohorts showed about <2, 40, and ~99% mortality, respectively, in response to storage and aging.
- A total of 19,340 genes were significantly differentially expressed (SDE) in imbibed axes compared to dry axes. Gene expression patterns of imbibed axes clustered into three groups that represented high, low, and no germination potential (GP). There were 17,360 SDE genes in high-GP axes and 4,892 SDE genes, mostly upregulated, in no-GP axes. Transcriptomes of no-GP axes were similar to healthy axes at 3 HAI.
- Slow transcription, not transcription errors or novel expression pathways, portends failure to transition from seed to seedling. We conclude that the signature of death in dry aged seeds arises from metabolism that is “too little and too late.”

## Introduction

Injuries accumulate in organisms dying of “old age.” Oxidative reactions, often involving reactive oxygen species (ROS), are implicated in aging (recently reviewed for plant germplasm by Leprince *et al.*, 2017; Waterworth *et al.*, 2019; Nagel et al., 2019; Zinsmeister *et al.*, 2020; Ballesteros *et al.*, 2020; Zhou *et al.*, 2020). In dry seeds, aging reactions affect multiple cellular components (Smith & Berjak, 1995; Walters, 1998; Rajjou *et al.*, 2008; Terskikh *et al.*, 2008; Kranner *et al.*, 2011; Colville *et al.*, 2012; El-Maarouf-Bouteau *et al.*, 2013; Michalak *et al.*, 2013; Kalemba & Pukacka, 2014; Xin *et al.*, 2014; Morscher *et al.*, 2015; Nguyen *et al.*,2015; Veselova *et al.*, 2015; de Souza-Vidigal *et al.*, 2016; Mira *et al.*, 2016, 2019, 2020; Nagel et al., 2019; Fleming *et al.*, 2017, 2018, 2019; Yin *et al.*, 2017; Kurek *et al.*, 2019; Wiebach *et al.*, 2019; Zhao *et al.*, 2020). Sometimes aging to death is attributed to a specific chemical change, such as oxidation of acyl double bonds in membrane lipids (Wiebach *et al.*, 2019; Sattler et al., 2004) or accumulation of abnormal L-isoaspartyl residues in the proteome of dry seeds (Ogé *et al.*, 2008). However, cause-effect relationships between change of *specific* cellular constituents and death have always been elusive, especially as we continue to discover more cellular components damaged by time. Accordingly, we have suggested that aging results from an accumulation of small changes, and at some point, a minor change has a major lethal effect, analogous to a ‘straw that breaks the camel’s back’. The distinction between mechanisms – mortality caused either by a specific major event or by many non-specific minor events – affects how we might detect aging, predict longevity and reinvigorate (if possible) damaged germplasm.

Deciphering the aging mechanism(s) of dry seeds is further complicated by our inability to pinpoint the time of death. There is a discrete time-frame during storage when dry germplasm has and then loses the potential to complete germination (Walters, 1998). The timing of this transition between alive and dead (i.e., longevity) varies considerably within and among species and cell types for reasons yet unknown (recently reviewed by Colville & Pritchard, 2019; Ballesteros *et al.*, 2020; Solberg *et al.*, 2020). Moreover, this transition occurs discreetly in dry seeds because we have limited tools to measure aliveness in “cryptobiotic” organisms. Therefore, we learn that a dry organism perished when it does not revive upon hydration. Did hydration deliver the lethal blow? Early research suggesting that hydration induces aging reaction cascades (Smith & Berjak, 1995) is supported by evidence of programmed cell death metabolism in aged, imbibing seeds (Kranner *et al.*, 2006; El-Maarouf-Bouteau *et al.*, 2011; Chen *et al.*, 2013; Wang *et al.*, 2015). Alternatively, early imbibition is a time for repairing damaged cellular components (Ogé *et al.*, 2008; Waterworth *et al.*, 2010, 2016, 2019; Rajjou *et al.*, 2012; Sano *et al.*, 2016) or activating transcription, translation and post-translational modifications essential for germination (Rajjou *et al.*, 2012; Galland *et al.*, 014; Bai *et al.*, 2017; Yin *et al.*, 2018; Sano *et al.*, 2020; Zhou *et al.*, 2020). Conceivably, death may occur when repair pathways take precedence over seedling development pathways (Masubelele *et al.*, 2005; Rosental *et al.*, 2014; Dirk & Downie, 2018; Cai, *et al.*, 2020; Sano *et al.*, 2020).

This paper focuses on the transcriptional machinery in dry-stored seeds as an essential component of germination potential (Rajjou *et al.*, 2012; Bai *et al.*, 2017; Dirk & Downie, 2018). Our study assesses the transcriptome of aged, *imbibed* seeds, compared to our earlier studies where we examined the transcriptome of aged, *dry* seeds. Numerous studies show continuous, possibly linear, degradation of RNA with time in dry storage (El-Maarouf-Bouteau *et al.*, 2013; Fleming *et al.*, 2017, 2018, 2019; Walters *et al.*, 2020; Sano *et al.*, 2020; Zhao *et al.*, 2020), which links rates of RNA degradation and longevity. However, correlation of mortality with degradation of a specific amount of total RNA or mRNA, or a specific transcript, was not demonstrated for dry seeds. We, therefore, explore a possible “death signature” for aged seeds during hydration. In particular, we looked for up- or down- regulation of a total of 3,492 ‘germination genes’. The subset of genes with changed regulation included transcription factors implicated in soybean seed longevity, enzymes critical for DNA repair (including PIMT, PARP, DNA ligases, and cyclins), homologs of genes required for germination in Arabidopsis (ABI3, HSFA9, PEPCK, and TIM), endo-beta-mannanases, LEAs, and enzymes involved in fermentation, lipid degradation, sugar metabolism, protein biosynthesis, and protein homeostasis (reviewed by Beilleny-Rabelo *et al.*, 2016; Pereira Lima *et al.*, 2017; Cai, *et al.*, 2020; Sano *et al.*, 2020; Zinsmeister *et al.*, 2020; Zhou *et al.*, 2020; also references in this paper).

We hypothesized that imminent death would be reflected by differences in expression patterns for key genes among healthy and dying seeds. We predicted down-regulation of key transcripts in dying seeds because the transcripts might be preferentially fragmented during dry storage (Fleming *et al.*, 2017, 2018, 2019) and further degraded during imbibition. We also postulated that *de novo* transcription, that might serve to replace damaged transcripts, would likely be impaired by damaged transcriptional machinery.

Radicle emergence occurs in healthy soybean seeds at about 24 hours after imbibition (HAI) and transcription patterns during early stages of normal germination are established (Bellieny-Rabelo *et al.*, 2016). We contrasted these established patterns with those of dying and recently dead soybean seeds from a unique collection of Williams ’82 cohorts harvested and stored since 1988 (Fleming *et al.*, 2017; Walters *et al.*, 2020). We compared transcriptomes among cohorts that were mostly vigorous or mostly dead, as well as a cohort exhibiting signs of rapid viability loss. A cotyledon greening assay was used to distinguish germination capacity in the dying cohort, which had nearly equal proportions of viable and inviable seeds. Instead of the predicted down-regulation of key transcripts, we found strong evidence that axes which could not complete germination still actively transcribed ‘germination genes’ but at a much slower pace.

## Materials and Methods

### Plant material

Soybean (*Glycine max* (L.) Merr, cv. ‘Williams 82’) seeds are part of a legacy collection of cohorts harvested between 1988 and 2019 (Fleming *et al.*, 2017). Seeds were received 3-6 months after harvest and germination percentages were high (between 98-100%) for all cohorts. Seeds were stored at 5 °C and approximately 30-50% relative humidity (RH). Under these conditions, germination percentages tend to decline after about 10 to 15 years (Walters *et al.*, 2020). For this study, we selected cohorts that either showed no evidence of aging (harvested in 2015, 2015H), severe aging (harvested in 1996, 1996H) and rapid viability loss (harvested in 1999, 199H). About half of the 1996H cohort was dead by 2008 (P50 = 12 years) and nearly all were dead by 2013 (data not presented). The 1999H cohort began exhibiting symptoms of aging in 2014 and 2015 assays (data not presented).

### Viability assessments

Seed cohorts were monitored for viability annually or biennially using germination assays. Germination assays occurred in 2017, 2018 and 2019, and flanked the time that RNA was extracted in 2018. Monitor tests consisted of about 50 to 200 seeds that were prehydrated overnight at near 100% RH, then rolled in wet paper towels (Anchor Paper Co., St Paul, MN, USA) and incubated in the dark at 25 °C for seven days. Seedlings were scored for normal germination following AOSA criteria (AOSA, 2012); radicle length was also measured to assess vigor.

We associated cotyledon greening with embryonic axis growth in all cohorts, especially in 1999H seeds. The reliability of this method was tested in a number of cohorts. Embryonic axes were separated from cotyledons of both dry seeds and seeds imbibed in the dark for 24 hours. Excised axes were placed on Murishage and Skoog basal medium (to visualize effects of surface sterilization and nutrients) or on dampened paper. Viable axes germinated in culture or expanded at least 2 cm on paper within 2-3 days after imbibition. Cotyledons were imbibed on paper for 24 hours in darkness, then placed in transparent plastic boxes in room light for 3-4 days and scored for whether they remained yellowish white, greened on the flat cotyledon surface or greened throughout. Thorough cotyledon greening invariably occurred in recently-harvested cohorts in which axes readily germinated (on nutrient medium) or expanded (on paper). Cotyledon greening rarely occurred in low-germinating cohorts stored for at least 22 years. In cohorts that had a mixed population of germinable and not-germinable seeds (such as 1999H), cotyledon greening provided a reliable marker of which axes were able to expand.

To compare germination speed among cohorts, time courses for water uptake, axis growth, and cotyledon greening were developed. Seeds rolled in wet paper towels (20-25 rolls containing 25-30 seeds each) were sampled every 2-8 hours for radicle emergence, cotyledon and axis fresh mass, and axis dry mass; dry mass was measured after heating axes at 95 °C for 2 days. Seeds were considered germinated when radicles elongated > 2 mm from the testa.

Cotyledons that were severed from axes at each time point were placed in light and observed periodically for color changes. For each roll, germination proportions were calculated as the number of germinated seeds divided by the number of seeds in the roll.

### RNA extraction, characterization, and sequencing

In 2018, we extracted RNA from embryonic axes for transcriptome sequencing. Seeds were imbibed at 25 °C in the dark and embryonic axes were excised at the cotyledonary node 24 hours after imbibition (HAI), leaving the plumule attached to the cotyledon. A total of 20 axes were excised from 1996H and 2015H seeds, and 60 axes were excised from 1999H seeds. Each axis was flash-frozen in liquid nitrogen and stored at −80C. To estimate germination potential, cotyledons were kept moist in constant room-lit conditions (cool white fluorescent lights). At 72 HAI (48 hours of cotyledon light exposure), each cotyledon was examined for greening, presence of microbial contamination, and plumule elongation. Based on this response, each excised axis was categorized as viable (1999H and all 2015H) or inviable (1999H and all 1996H).

Individual embryonic axes, excised from dry (0 HAI) and hydrated (24 HAI) seeds, were ground to a fine powder in a Retsch (Haan, Germany) Bead Mill under liquid nitrogen in microcentrifuge tubes containing 1 mg of polyvinylpyrrolidone-40 (Fisher Scientific, Fair Lawn, NJ, USA). RNA was extracted from each ground axis using the Qiagen (Hilden, Germany) Plant RNeasy kit following the recommended protocol, with the additional step of repeating the final wash with 500 μL of buffer RPE to reduce guanidine hydrochloride contamination.

RNA yield was quantified using a Nanodrop 1000 spectrophotometer (Thermo Fisher Scientific, Wilmington, DE, USA). Samples were diluted to 1 ng μl^−1^ of nucleic acids in nuclease-free water. Integrity of diluted RNA was quantified on an Agilent (Waldbronn, Germany) Bioanalyzer, using Agilent RNA 6000 Pico chips and the Plant RNA Pico assay (Agilent 2100 Expert software version B.02.08.SI648 R3), following the manufacturer’s protocols. Five axes producing the highest RNA Integrity Numbers (RINs) for each combination of cohort (1996H, 1999H, 2015H), imbibition time (0 or 24 HAI), and viability category (for 1999H imbibed axes), were selected for sequencing.

Total RNA was submitted to the University of Delaware Sequencing and Genotyping Center. After poly-A selection, 1 μL of a 1:50 dilution of ERCC Spike-In mix (Ambion, Thermo Fisher Scientific, Wilmington, DE, USA) was added to each sample. All libraries were pooled and sequenced on two lanes of an Illumina (San Diego, CA, USA) HiSeq with 250 bp paired-end reads. The sequencing dataset for Williams ’82 axes from 1996H, 1999H, and 2015H cohorts is subsequently referred to as the “storage time experiment”.

Transcriptome data from the storage time experiment were compared with RNA-seq data collected for healthy soybean axes of cv. ‘BRS 284’ at 0, 3, 6, 12, and 24 HAI (Bellieny-Rabelo *et al.*, 2016). All 50 datasets from this experiment were downloaded from the Sequence Read Archive using SRA Explorer (sra-explorer.info) and project accession number PRJNA326110. This sequencing dataset is subsequently referred to as the “imbibition time experiment”. All analyses were conducted on both the storage time and imbibition time experiments simultaneously.

### Identifying significantly differentially expressed transcripts

Adapters were trimmed from all reads with CutAdapt v. 2.5 (Martin, 2011), and any reads < 15 bp were excluded. Reads were aligned to the most recent soybean reference transcriptome (Gmax_508_Wm82.a4.v1.transcript.fa; Joint Genome Institute, Phytozome) using hisat2 v. 2.1.0 (Kim *et al.*, 2019) with standard parameters, and converted to indexed bam files using SAMtools v. 1.9 (Li *et al.*, 2009). The number of reads associated with each transcript (counts) was extracted from indexed bam files using the SAMtools function ‘idxstats’ (Table **S1,S2**). Differential expression of transcripts in imbibed versus dry samples was identified in R (R Core Team, 2018) using edgeR v. 3.26.8 (Robinson *et al.*, 2010; McCarthy *et al.*, 2012).

Imbibed samples were compared to the dry controls from their respective experiment (storage time or imbibition time). Before calculating differential expression, lowly-expressed transcripts were automatically identified and removed from further consideration using the filterByExpr function, and libraries were normalized based on library size using the calcNormFactors function. Transcripts were considered significantly differentially expressed (SDE) when |log_2_ fold-change (FC) in expression| ≥ 2 and adjusted p-value < 0.001 (Table **S3,S4**).

### Functional characterization and transcriptome clustering

An X4 MapMan v. 3.6.0RC1 (Thimm *et al.*, 2004) mapping for soybean was generated with Mercator4 v.2 (Schwacke *et al.*, 2019). Transcripts were automatically assigned to annotation bins (Table **S5**), which were then used to assess over-representation of annotations within similarly-expressed clusters of transcripts (see below). Log_2_FC for transcripts SDE in a given treatment were visualized using the ‘X4.1 Metabolism overview R1.0.xml’ map.

Heatmaps were generated in R using pheatmap v. 1.0.12 (Kolde, 2019) to cluster experimental treatments and transcripts by expression values. Log_2_FC values were converted to z-scores [z = (x – μ)/**σ**, where x is the observed value, μ is the sample mean and **σ** is the sample standard deviation] before plotting, confining values to the same scale for each transcript. A negative z-score indicates that log_2_FC for that transcript in that sample was lower than the mean across all samples. As the imbibition time experiment used much deeper coverage than the storage time experiment, z-score transformation was done independently for each experiment.

Any transcript identified as SDE in the storage time experiment was included in the heatmap (19,340 transcripts). Transcripts were clustered using z-score transformed log_2_FC values from the storage time experiment, calculating the distance matrix with Euclidean distance and using the complete linkage method for hierarchical clustering (Table **S6**). Experimental groups (no-, low-, and high-germination potential 24 HAI [storage time experiment] as well as 3, 6, 12, and 24 HAI [imbibition time experiment]) were clustered with the same clustering parameters, using z-score transformed values from all experiments.

After clustering, a two-proportion z-test was performed to identify which MapMan annotation categories were over- or under-represented in each cluster of transcripts. This test compares the abundance of a category within a cluster to the abundance of that category in the entire population of SDE transcripts. A Bonferroni-corrected alpha level of 0.0003 (0.05/174) was used to determine categories with significant over- or under-representation within a cluster (Table **S7**).

### ‘Germination genes’

A literature survey was performed to identify genes important for germination. Most genes were characterized in *Arabidopsis*; *Glycine max* homologs to *Arabidopsis* genes were identified using TAIR (arabidopsis.org). In Soybase (soybase.org/correspondence), gene IDs were converted from genome version a2 to a4 and paralogous genes within the *Glycine max* genome were identified. In some cases, only a general class was identified (*e.g.*, “fermentation”), which could always be matched to a MapMan annotation bin. Expression of all genes within the bin was examined (Table **S8**). The expression of 5,712 transcripts (corresponding to 3,492 ‘germination genes’) that were SDE in any GP category was visualized with pheatmap using the same z-score adjustment and clustering parameters described above (Table **S9**).

## Results

### Seed health

#### Germination time course and cotyledon greening assays

Germination percentages, measured four times between 2017 and 2019, were 98-99% (2015H), 60-90% (1999H) and 0-3% (1996H). A germination time course, conducted in 2019, depicts germination events which indicate health of the cohort as well as individual seeds. Radicle emergence was first observed 22 HAI in the 2015H and 1999H cohorts, though at different proportions (9 and 1%, respectively) (Fig. **1a**). Within 36 HAI, radicles emerged from 95% and 42% of seeds from the 2015H and 1999H cohorts, respectively, compared to 0% in the 1996H cohort. By 72 HAI, total germination of 95, 64 and 1% was observed for 2015H, 1999H and 1996H cohorts, respectively. Water content increased from 0.076 - 0.096 g H_2_O g^−1^ dry weight (dw) for dry embryonic axes to 1.2 - 1.4 g H_2_O g^−1^ dw 12 HAI for all cohorts (Fig. **1b**). Rapid increases in water uptake and fresh mass occurred at 32 and 42 HAI for 2015H and 1999H axes, respectively, and was not observed for 1996H axes (Fig. **1b,c**). Axes from the 2015H cohort began to accumulate dry matter 32 HAI, but this capacity was weakly or not observed in 1999H or 1996H axes even 66 HAI (Fig. **1d**).

**Figure 1.**
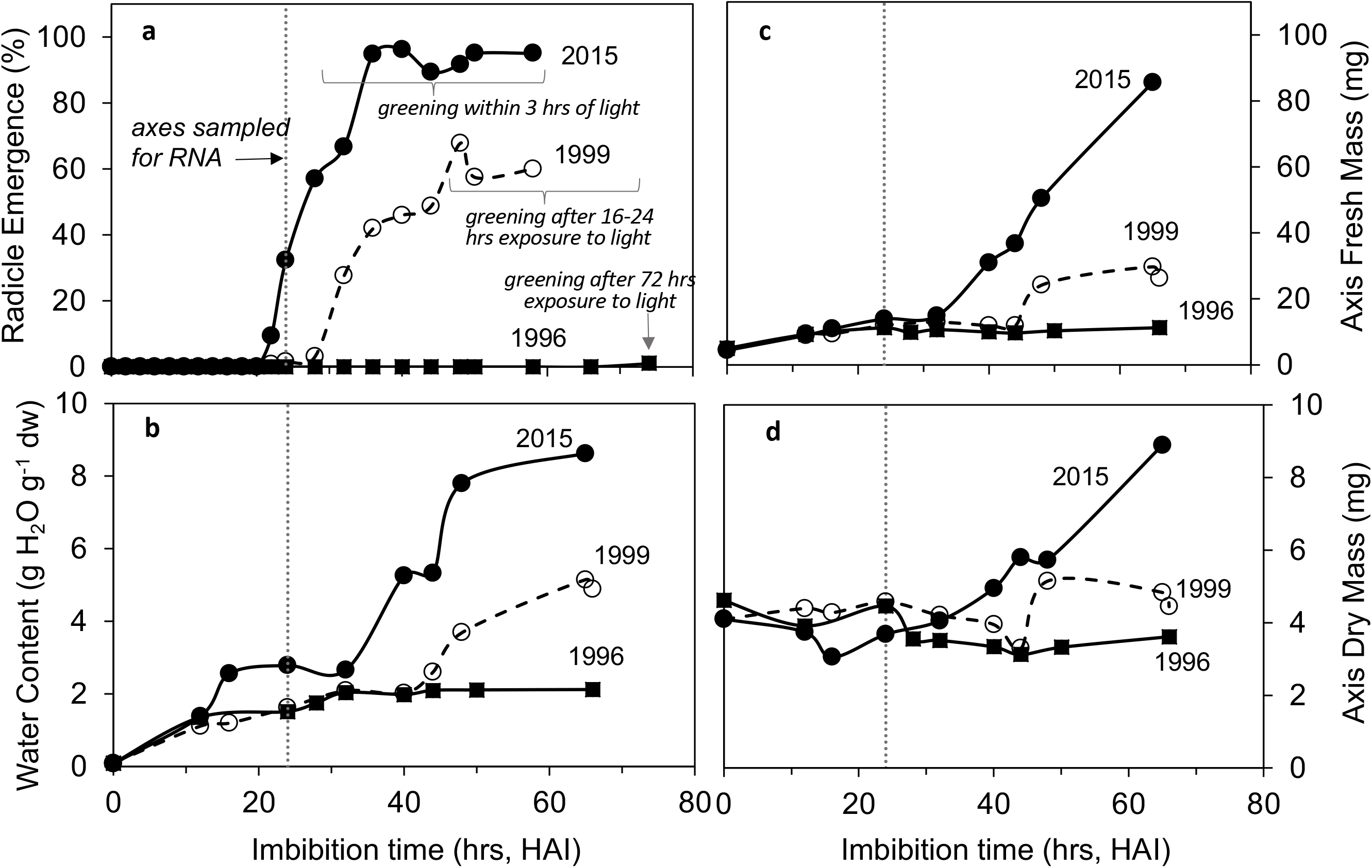
Markers of germination capacity in imbibing soybean (cv. ‘Williams 82’) seeds harvested in 2015 (filled circles), 1999 (open circles, dashed curve) and 1996 (filled squares). Data reflect % radicle emergence (**a**), axis water content (**b**), and axis fresh (**c**) and dry (**d**) mass measured at indicated times for separate damp paper towel rolls containing 20-30 seeds incubated at 25C. The dotted vertical line indicates timing of RNA extraction of isolated axes and light exposure of corresponding cotyledons.

The cotyledon greening assay reliably indicated axes that would and would not elongate and cotyledon attachment was not required for axis expansion or cotyledon greening (**Fig. S1**). Greening was rapid in 2015H seeds, usually occurring after 3 hours exposure to light (data not shown). Greening took longer in 1999H seeds (data not shown) and rarely occurred in 1996H seeds.

#### RNA integrity among seed cohorts

RNA was extracted in 2018 from dry and imbibed embryonic axes, providing data for seeds stored for 3, 19 and 22 years. Notably, RNA Integrity Numbers (RINs) neither increased nor decreased in imbibed samples compared to dry samples of the same cohort (*p* > 0.05, Tukey’s HSD, Fig. **2**, Table **S10**). RINs were similar for 2015H and 1999H axes, regardless of whether their corresponding cotyledons turned green (*p* > 0.05, Tukey’s HSD). RINs were significantly lower for 1996H axes compared to the 1999H and 2015H cohorts (*p* < 0.05, Tukey’s HSD) (Fig. **2**).

**Figure 2.**
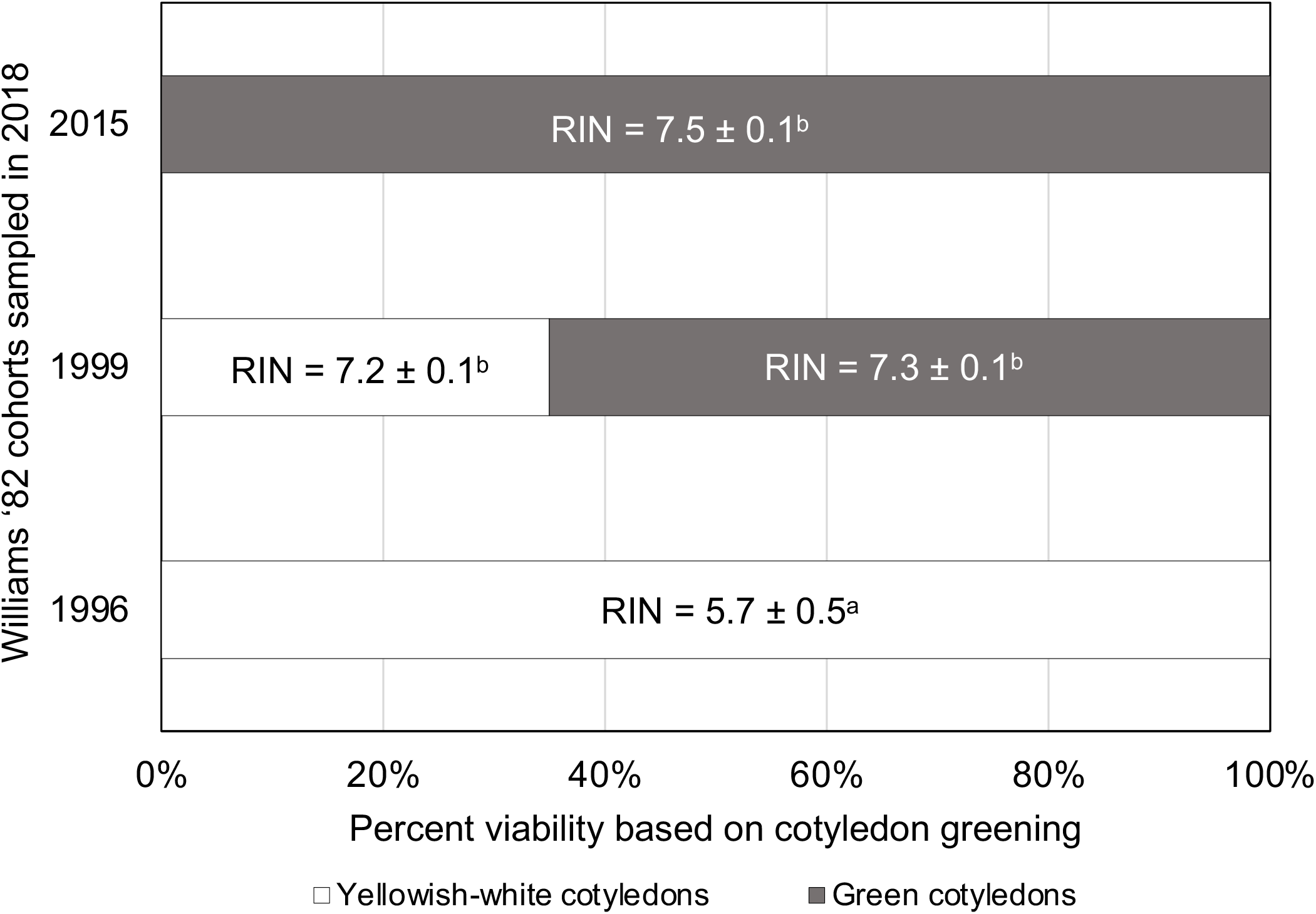
RNA integrity, as measured by RIN (Schroeder *et al.*, 2006) of five excised embryonic axes from different soybean (cv. ‘Williams 82’) cohorts imbibed for 24 hours. Shaded boxes represent the proportion of 100, 200, and 100 axes for 2015H, 1999H and 1996H cohorts, respectively, in which corresponding cotyledons greened (indicating positive germination capacity of axes). Values in each bar represent the average RIN ± std deviation for each cohort and viability class. RINs marked with different superscript letters are significantly different at p < 0.05) (see Table **S2** for details).

Consistent with the RINs, electropherograms from 2015H and 1999H axes 24 HAI were mostly indistinguishable, showing typical patterns of high-integrity plant RNA, with two prominent peaks for the 18S and 25S rRNA subunits, minor peaks for the 5S rRNA subunit and plastid rRNAs, and an absence of prominent peaks in the “fast” region (27.25-41 s) (**Fig. S2**).

Electropherograms from 1996H dry and imbibed axes had reduced 25S peaks relative to 18S peaks, as well as more prominent peaks in the “fast” region, indicating RNA fragmentation as reported previously (**Fig. S3**) (Fleming *et al.*, 2017, 2018. 2019).

#### Transcriptome overview

Global differences in log_2_FC expression among transcriptome libraries from individual embryonic axes were compared in a multidimensional scaling plot (Fig. **3**). Instead of clustering by harvest year or cotyledon status, axes separated into four clusters corresponding to dry axes (all cohorts), axes from seeds with green cotyledons [“high germination potential” (GP), 1999H and 2015H], axes from seeds with white cotyledons (“no-GP,” 1996H and 1999H), and a fourth group including axes from both 2015H and 1999H seeds having white or green cotyledons. This intermediate cluster, named “low-GP”, was spatially separated from the two other 24 HAI axis clusters and was treated separately in subsequent analyses.

**Figure 3.**
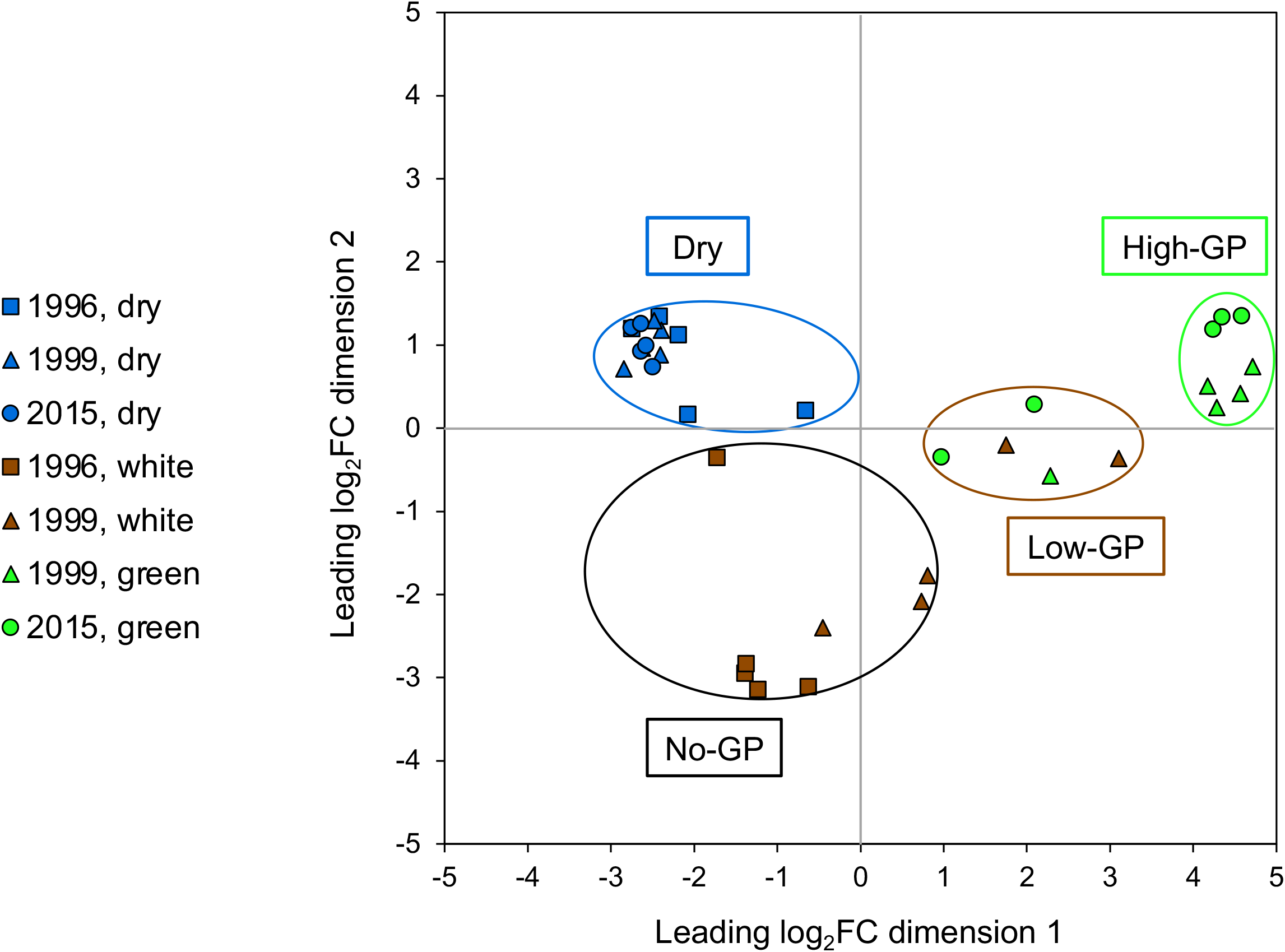
A multi-dimensional scaling (MDS) plot summarizing relationships between transcriptomes, where each point represents the transcriptome from a single dry (blue symbols) or imbibed (green or brown symbols) soybean (cv. ‘Williams 82’) embryonic axis, and proximity indicates higher similarity. Imbibed axes have corresponding cotyledons that did or did not green, indicated by green and brown symbols, respectively. Seed cohort is represented by circles (2015H), triangles (1999H) and squares (1996H). Axes clearly separated based on dry/imbibed. Imbibed axes sorted into three groups based on cotyledon color, interpreted as representing the entire seed’s likely germination potential (GP): all white cotyledons (no-GP), all green cotyledons (high-GP), and a mixture of white and green cotyledons (low-GP).

Overall, in the storage time experiment, 19,340 genes were significantly differentially expressed (SDE) in imbibed compared to dry axes, according to stringent criteria of |log_2_FC| > 2 and p-value < 0.001. More than half (58%) of SDE genes were shared among GP categories (Fig. **4**, Table **S3**). High-GP axes differed most from dry axes (17,360 SDE genes) and no-GP axes differed least from dry axes (4,892 SDE genes). Most (94%) of the SDE genes in low-GP axes were also SDE in high-GP axes.

**Figure 4.**
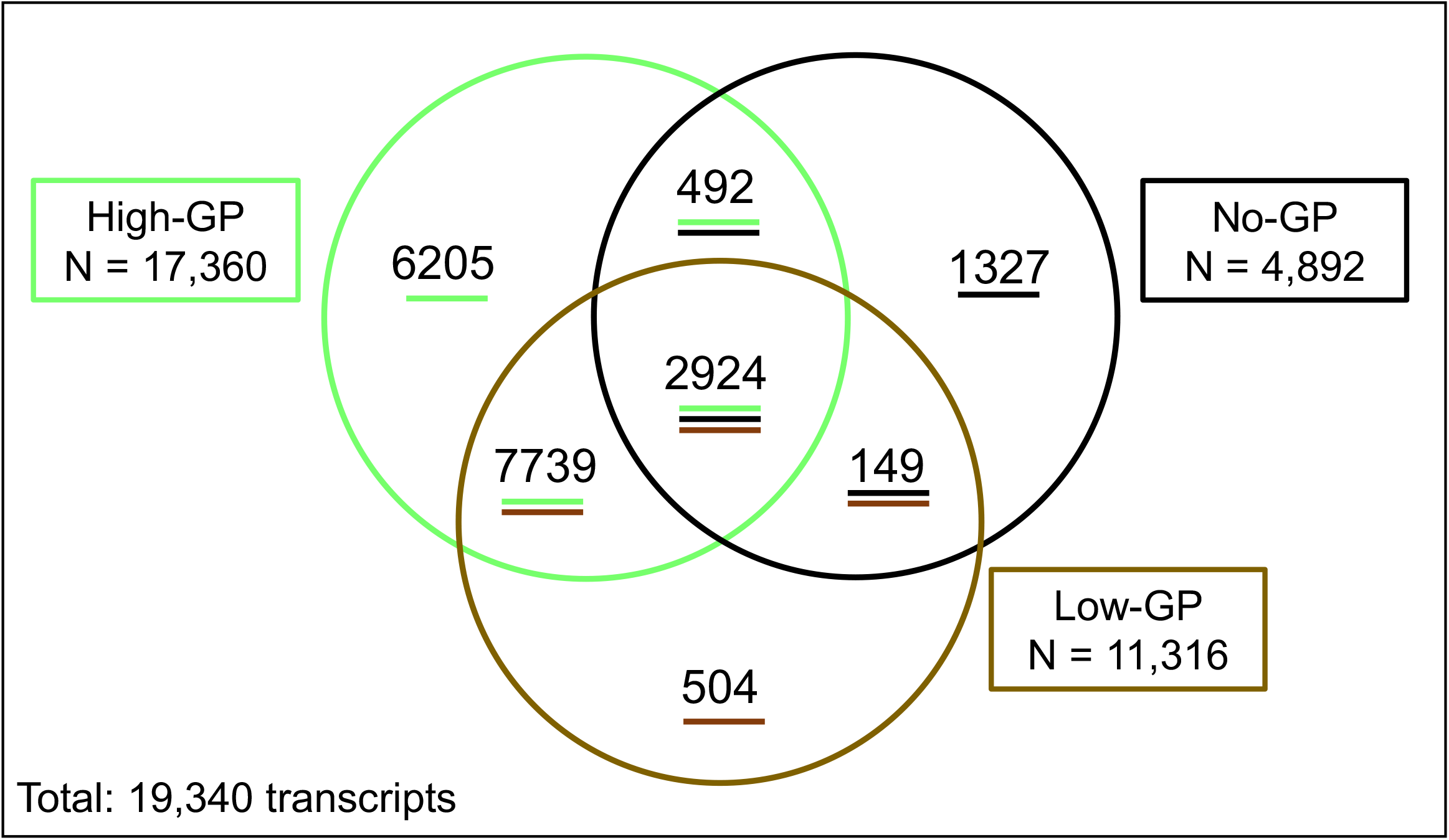
Venn diagram of transcripts significantly (|log_2_FC| > 2 and p < 0.001) differentially expressed in soybean (cv. ‘Williams 82’) axes from different cohorts 24 hours after imbibition compared to dry axes from all cohorts. A total of 19,340 transcripts were differentially expressed in at least one germination potential (GP) category, indicated by clusters in Fig. **3**. High-GP axes had the largest number of differentially expressed transcripts, and no-GP axes had the fewest. The majority of transcripts differentially expressed in no-GP axes were identified in all GP categories. Low-GP axes had an intermediate number of differentially expressed transcripts, most of which were shared among high- and no-GP axes.

Of the genes SDE in each GP category, 67, 76 and 90% of transcripts were upregulated in high-, low- and no-GP axes, respectively (see upper quadrants in Fig. **5a,b,c**). Gene expression in high- and low-GP axes was highly correlated (R^2^ = 0.94, P << 0.01), and the slope of correlation, 1.27 ± 0.01 (uncertainty of slope at 95% confidence) indicated more intense DE in high-compared to low-GP axes (Fig. **5a**). The correlation was weaker between high-GP and no-GP axes (R^2^ = 0.57, P << 0.01, slope = 1.25 ± 0.04 at 95% confidence), but the slopes were not significantly different (t-test of slopes not significant at P > 0.15) (Fig. **5b**). Some transcripts were SDE in both high- and no-GP axes, but in opposite directions (lower right and upper left quadrants). Comparisons of the log_2_FC for low- and no-GP axes were also significantly correlated (R^2^ = 0.69, P << 0.001), and the slope of 1.06 ± 0.03 was significantly different than correlations involving the high-GP group (t-test of slopes significant at P << 0.01) (Fig. **5c**).

**Figure 5.**
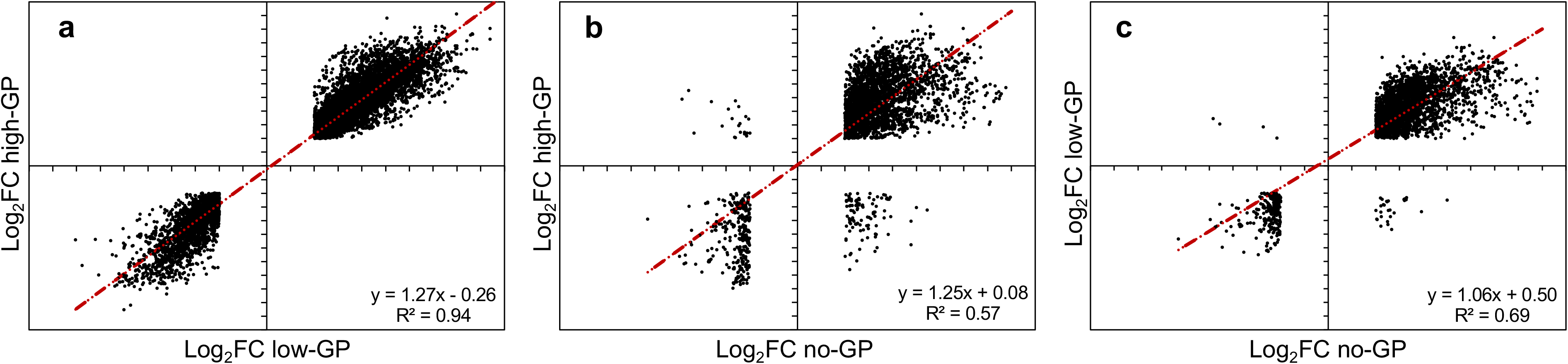
Correlations between intensity of expression (log_2_FC) of significantly differentially expressed soybean genes shared between different germination potential (GP) categories: 10,663 genes shared between high- and low-GP (**a**), 3416 genes shared between high- and no-GP (**b**) and 3073 genes shared between low- and no-GP axes (**c**). The equation for the regression line and the correlation coefficient for each relationship are indicated. All relationships are significant at P << 0.01. Slopes between regression lines in (**a**) and (**b**) are not significantly different (P > 0.05), but the slope of the regression line in (**c**) is significantly different from the slopes in (**a**) and (**b**) (P < 0.05). Of note are the number of genes in the upper left and lower right quadrants in (**b**) and (**c**), indicating opposite regulation of these genes in no-GP axes that lowers the correlation coefficient.

#### Functional differences in transcriptomes of aged versus healthy seeds

Transcripts were assigned to MapMan annotation categories, and log_2_FC values for SDE genes in each GP category were mapped based on their annotation to understand how metabolism might differ between GP categories (Fig. **6**, Tables **S3,S5**). High-GP axes had SDE genes distributed among all major metabolic pathways (Fig. **6a**), including pathways essential for photosynthesis (light reactions, ROS, Calvin cycle). Other categories with many SDE genes in high-GP axes included catabolism of lipids and raffinose oligosaccharides, compounds which accumulate in the embryonic axis during soybean seed development. MapMan annotation categories were more sparsely populated in low- and no-GP axes because there were fewer SDE genes (Fig. **6b,c**). For example, genes involved in lipid or carbohydrate mobilization or nucleotide metabolism are somewhat and barely represented in low- and no-GP axes, respectively. This overview of embryonic axis metabolism 24 HAI revealed no dominant metabolic category for any GP class. SDE genes in no-GP axes, which had 72% less differential expression than high-GP axes, were distributed among annotation categories in approximately the same ratio as in high-GP axes.

**Figure 6.**
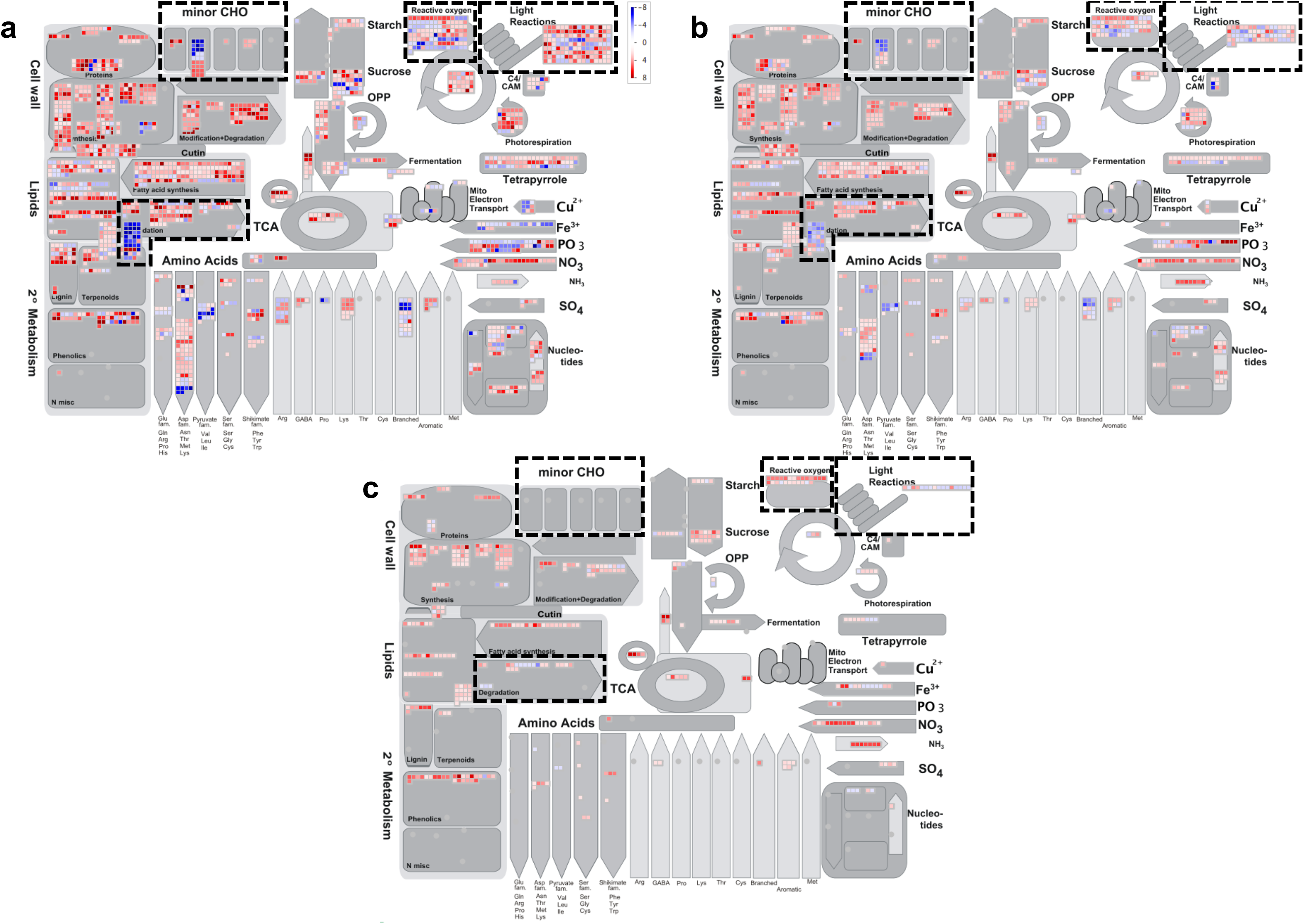
Mapman displays of genes significantly differentially expressed (SDE) in high germination potential (GP) (**a**), low-GP (**b**) and no-GP (**c**) soybean (cv. ‘Williams 82’) axes. Various gray shapes represent metabolism categories and red and blue squares represent up- and down-regulation of SDE genes, respectively. Intensity of color represents |log_2_FC| with light pink or blue indicating |log_2_FC| = 2 and most intense color indicating |log_2_FC| = 8. High-GP axes (**a**) show high |log_2_FC| values in several metabolism categories that is absent from no-GP axes (**c**) and intermediate in low-GP axes (**b**). Metabolism categories discussed in the text are noted with a dashed outline.

#### Gene expression during imbibition: from dry seed to completed germination or death

Gene expression patterns of healthy soybean seeds during an imbibition time course provide a useful context for interpreting the effects of storage time. Transcriptomic data from an imbibition time course of healthy soybean (cv. ‘BRS 284’) axes (“imbibition time experiment”, Bellieny-Rabelo *et al.*, 2016) were compared to transcriptomes of high-, low- and no-GP axes (“storage time experiment”). In the imbibition time experiment, the number of SDE (|log_2_FC| > 2, p-value < 0.001) genes in imbibed compared to dry (0 HAI) axes increased at each time point, with 2,888, 7,267, 14,796, and 28,032 SDE genes observed at 3, 6, 12 and 24 HAI (**Fig. S4**, Table **S4**). A core set of 1917 genes was SDE at all time points, and 982 of these genes were also SDE in high-, low- and no-GP axes. More genes were up-regulated than down-regulated at all imbibition time points, but the number of down-regulated genes increased from 1% to 33% of SDE genes between 3 and 24 HAI.

Similarly to the storage-time experiment, SDE transcripts were found in all the major metabolic pathways at all time points in the imbibition-time experiment. The magnitude of differential expression increased with imbibition time (Fig. **7**), and at 24 HAI, gene expression patterns for ‘BRS 284’ axes were similar to high-GP ‘Williams 82’ axes, with all metabolic pathways represented (compare Fig. **7d** with Fig. **6a**). In healthy ‘BRS 284’ axes 3 HAI, fewer genes were SDE, and these genes had smaller changes in differential expression (Fig. **7a**), compared to no-GP ‘Williams 82’ axes 24 HAI (Fig. **6c**).

**Figure 7.**
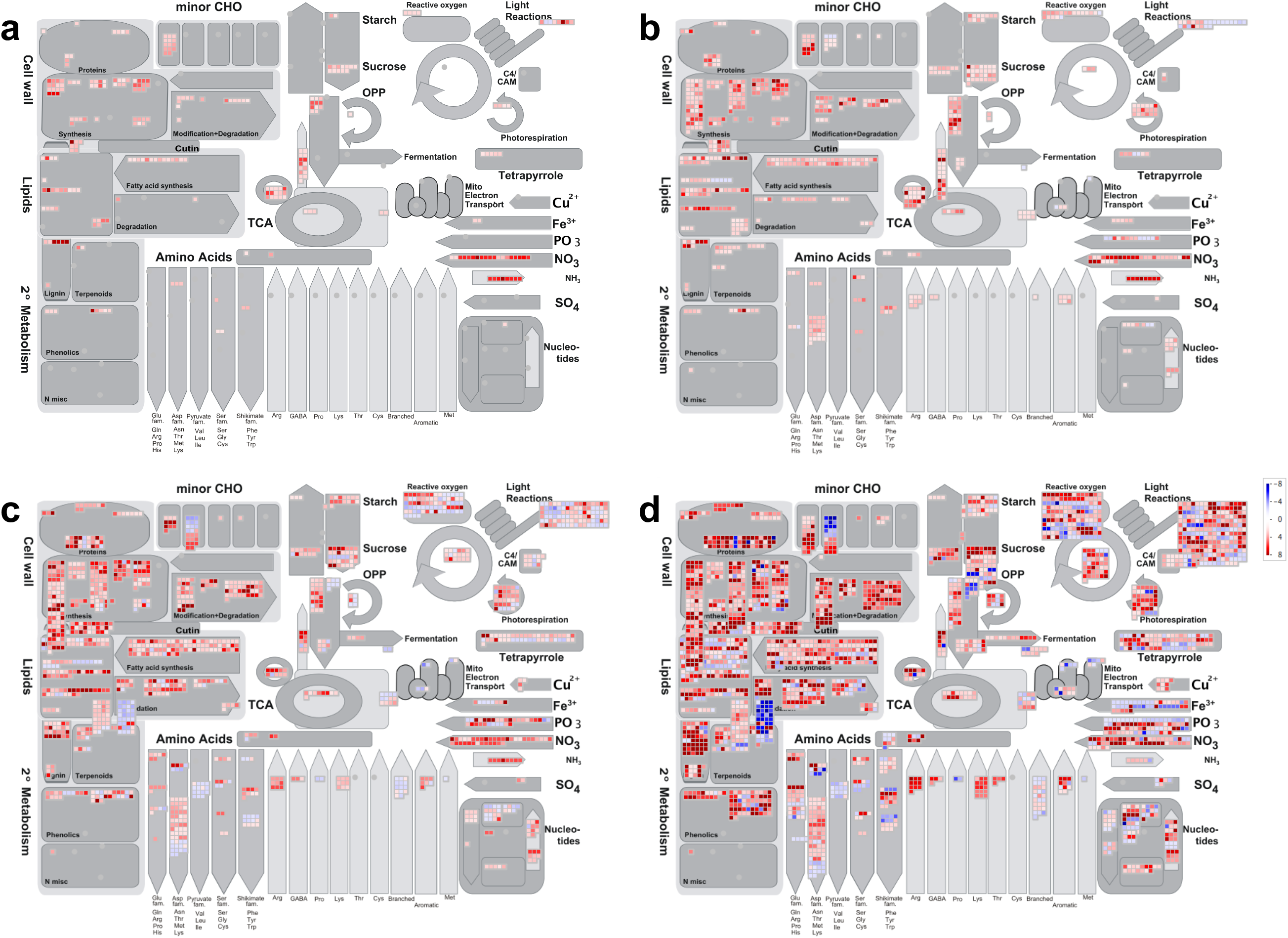
Mapman displays of genes significantly differentially expressed (SDE) in imbibed compared to dry soybean (cv. ‘BRS 284’) axes at sequential imbibition times based on data from Bellieny-Rabelo et al. (2016), illustrating increased expression at 3 hours after imbibition (HAI) (**a**), 6 HAI (**b**), 12 HAI (**c**) and 24 HAI (**d**). As with Fig. **6**, red and blue squares represent up- and down-regulation of SDE genes, respectively. Intensity of color represents |log_2_FC| between 2 and 8.

To further compare expression patterns between storage time (high-, low-, and no-GP in cv. ‘Williams ‘82’ axes 24 HAI) and imbibition time (3, 6, 12, and 24 HAI in healthy cv. ‘BRS 284’ axes), we used a heatmap of the 19,340 genes identified as SDE in any GP category in the storage-time experiment (Fig. **4**,**8**). Hierarchical clustering by treatment (GP category or imbibition time) showed that high-GP axes (Fig. **6a**) clustered with ‘BRS 284’ axes 24 HAI; low-GP axes (Fig. **6b**) clustered with ‘BRS 284’ axes 6-12 HAI; and no-GP axes (Fig. **6c**) clustered with ‘BRS 284’ axes 3 HAI. Axes of different GP categories did not cluster together, and only 6- and 12-HAI axes, among all imbibition time-points, clustered together.

**Figure 8.**
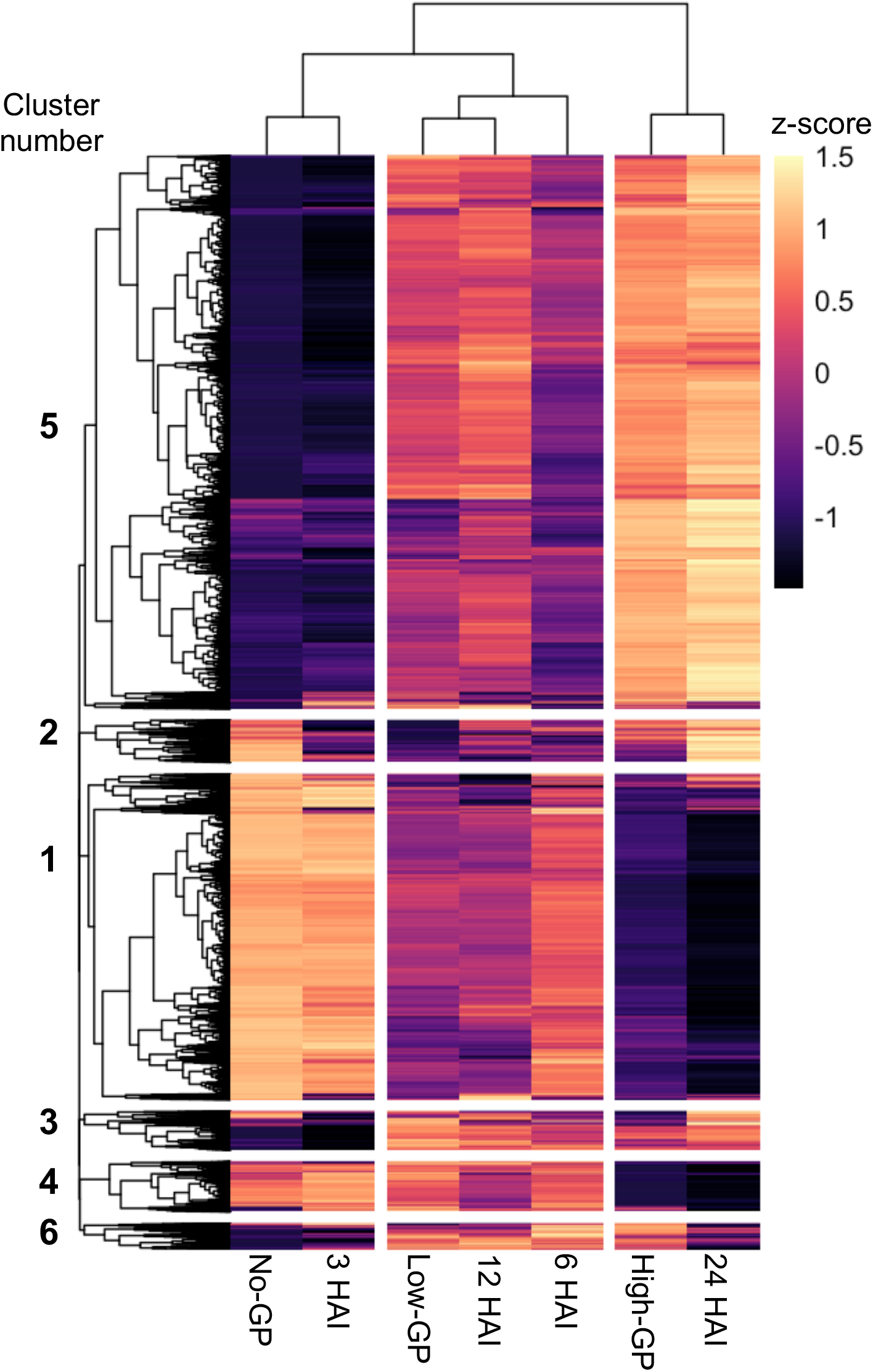
Heat map of all 19,340 transcripts significantly (|log_2_FC| > 2, p < 0.001) differentially expressed in any germination potential (GP) category (compared to dry soybean cv. Williams 82 axes) that were also expressed at all four imbibition time-points [3, 6, 12, and 24 hours after imbibition (HAI)]. Log_2_FC values were corrected by z-score independently for GP categories and imbibition time-points. Rows were clustered by expression profile in the three GP categories; columns were clustered by expression profile in all seven treatments. Six major clusters were observed among transcripts; three clusters were observed among treatments. No-, low- and high-GP axes clustered independently from each other; increasing GP was associated with longer imbibition times. Generally, intensity of expression was opposite in no-GP/3 HAI axes compared to high-GP/24 HAI axes, while expression in low-GP/6 HAI/12 HAI axes was intermediate.

Six general gene expression clusters were identified based on expression in high-, low-, and no-GP axes. The largest (Cluster 5: 10,325 genes) included genes with highest expression in high-GP and heathy axes 24 HAI and lowest expression in no-GP axes and healthy axes 3 HAI. A smaller cluster (Cluster 1: 6,073 genes) showed the opposite pattern, with highest expression in no-GP and healthy axes 3 HAI and lowest expression in high-GP and healthy axes 24 HAI. In most clusters, approximately average expression was found in low-GP axes as well as healthy axes 6 or 12 HAI (Table **S6**).

Genes belonging to a cluster may share functions as well as expression patterns. To identify functions associated with the different clusters, we tested whether any of the 28 MapMan annotation categories were significantly over- or under-represented in any cluster using a two-proportion z-test (Table **S6,S7**). Significantly over-represented categories in Cluster 5 (highest expression in high-GP/24 HAI axes) included cell cycle organization, cell wall organization, photosynthesis, and cytoskeleton organization; cytoskeleton organization was also over-represented in Cluster 2. Categories significantly under-represented in Cluster 5 included RNA biosynthesis and RNA processing. RNA processing, along with protein biosynthesis and homeostasis, were over-represented in Cluster 1 (highest expression in no-GP/3 HAI axes).

#### A closer look at genes important for seed germination in healthy and aged seeds

A total of 5,712 transcripts, representing 3,492 genes of known importance for seed germination, were identified from the literature. Only 27% of these “germination genes” were SDE in high-, low-, *or* no-GP axes from the storage time experiment (Table **S8**). Hierarchical clustering by treatment (GP category and imbibition time), using only the SDE germination genes (Fig. **9**, Table **S9**), produced identical clustering as in Figure 8, in which all SDE genes were considered. Most functional categories had a mixture of expression patterns, with some genes showing highest expression in no-GP/3HAI axes and lowest expression in high-GP/24 HAI axes, and the remaining genes showing the opposite expression pattern. However, all SDE ABI3 homologs (Category F, Fig. **9**), longevity-associated transcription factors (Category J) and mRNA quality control genes (Category Q) showed highest expression in no-GP/3 HAI axes, while all SDE PARPs (Category D) and fermentation genes (Category H) showed highest expression in high-GP/24 HAI axes.

**Figure 9.**
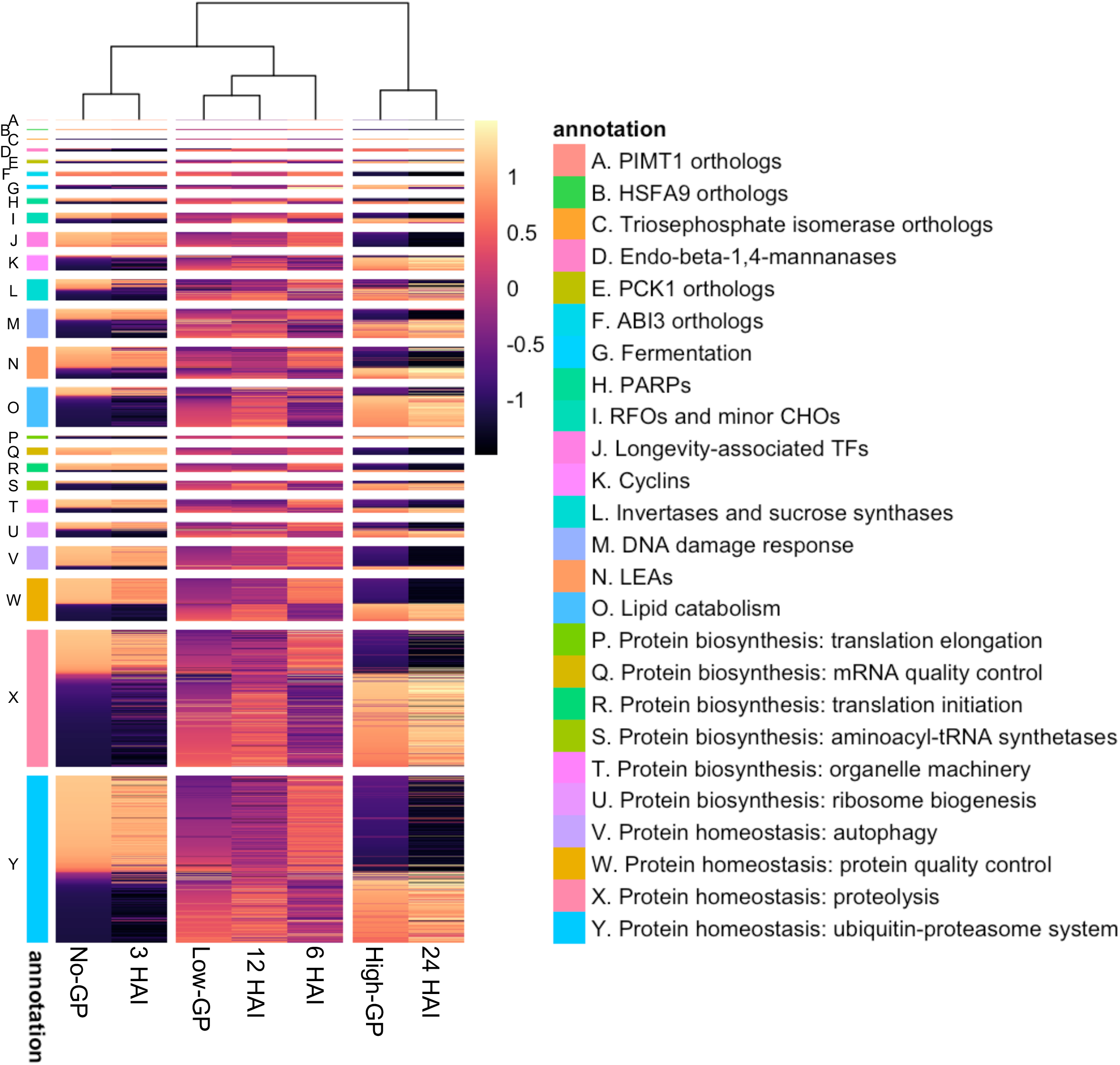
Heat map of z-score-corrected log_2_FC for 5712 transcripts involved in germination (see Supplemental Table 7), ordered on the y-axis by annotation and decreasing z-score in no-GP soybean axes, and clustered on the x-axis by expression profile. Log_2_FC values were corrected by z-score independently for GP categories and imbibition time-points. Genes were significantly differentially expressed in at least one of the GP categories. The same patterns observed in Fig. **8** were seen here also: GP categories did not cluster together, and increasing GP was associated with increased imbibition time; opposite expression intensity was found in no-GP/3 HAI versus high-GP/24 HAI, while low-GP/12 HAI/6 HAI axes were intermediate. No misregulation of germination genes in no- or low-GP categories compared to their early imbibition counterparts was apparent.

## Discussion

In this paper, we sought a transcriptional signal corresponding to lost viability in hydrated soybean seeds that had been stored dry for decades. We hypothesized that seeds become inviable during imbibition (Smith and Berjak, 1995), because transcriptomes of dry seeds did not reveal a “death” signal (Fleming *et al.*, 2017, 2018, 2019). We sequenced transcriptomes of dry and imbibed axes from three seed cohorts at different stages of degradation (Walters *et al.*, 2020): no loss in viability (98-99% germination, 2015H), almost complete loss (0-3% germination,1996H), and rapid loss in viability (60-90% germination depending on assay conditions, 1999H) (Walters *et al.*, 2020). Transcriptomes of healthy, dying, and dead dry axes were similar. Imbibition for 24 hours resulted in three distinct patterns of gene expression, which we named high-, low- and no-germination potential (GP) (Fig. **3**). Healthy axes (2015H) were found in both high- and low-GP categories; dying axes (1999H) were found in all three GP categories. Transcript expression was less intense overall, and few genes were down-regulated, in the no-GP cluster compared to the high-GP cluster (Fig. **5b**). However, comparing no-GP axes (imbibed for 24 hours) to healthy axes imbibed for less time showed that the expression profile of no-GP axes was highly similar to healthy axes imbibed for just 3 hours (Fig. **6c**,**7a,8**). These data contribute to the growing understanding of a metabolic burst that transitions dry seeds into seedlings (Rajjou et al., 2012; Galland et al., 2014; Bellieny-Rabelo et al., 2016; Bai et al., 2017). Our work further suggests that damage to seeds during dry storage decreases metabolic competence, reducing the metabolic burst to a fizzle.

### Imbibition allows significant up- and down-regulation of genes in vigorous axes

Earlier studies suggested that RNA fragments during dry storage, and we detected this tendency in mRNA using whole-molecule MinION sequencing techniques (Fleming *et al.*, 2018). The current study is based on Illumina sequencing datasets, which showed similar transcriptomes among dry ‘Williams 82’ axes stored for 3 to 22 years (Fig. **3**) even though they differed in viability (Fig. **1**) and RNA integrity (Fig. **2**). The Illumina platform provides sufficient coverage depth to compare differential expression across 35 samples, but this comes at the expense of observing RNA fragmentation.

Differential expression in high-GP ‘Williams 82’ axes and ‘BRS 284’ axes 24 HAI was similar (Fig. **8**). For example, genes involved in photosynthesis were found to be upregulated, indicating preparation for transitioning to autotrophic growth (Fig. **6**,**7**, Table **S6**). The similarity of transcriptomes of healthy axes from different cultivars allowed us to compare expression patterns between ‘Williams 82’ axes of different GPs and ‘BRS 284’ axes imbibed for different times.

Bellieny-Rabelo *et al.* (2016) identified many genes that were previously implicated in germination by comparing changes in adjacent time points during imbibition. Our goal, rather, was to uncover differences between alive and dead seeds that were fully imbibed and ready (or not) to complete germination. Many of the genes implicated in germination were not flagged in our study, perhaps because transient up- and down- regulation is completed by our sampling time at 24 HAI. Alternatively, identification of ‘germination genes’ during imbibition may be masked because their transcripts or proteins were already produced during seed maturation (Pereira Lima *et al.*, 2017), except for a few genes within broad metabolic categories; our analyses do not reveal candidates. Overall, our findings suggest that germination potential hinges on swiftly reaching a baseline level of metabolic competence, rather than effective transcription or translation of specific ‘germination genes.’

### Axes with no germination potential have functional transcriptional machinery

Transcriptomes of no-GP axes 24 HAI were distinct from dry axes and from healthy imbibed axes (Fig. **3**). That said, many SDE genes in no-GP axes that were shared by low- and high-GP axes were upregulated (Fig. **4**,**5**), even though the same genes were strongly down-regulated in high-GP axes (note low R^2^ for log_2_FC in high-GP versus no-GP axes in Fig. **5b**). In fact, no-GP axes produced new transcripts, although their weaker expression and misregulation in comparison to high-GP transcriptomes suggests that no-GP transcriptomes may be too small or uncoordinated to complete germination. We saw no evidence of divergent metabolic pathways in high-GP and no-GP axes that explain failed germination in the latter.

The transcriptome of no-GP axes 24 HAI largely reflects expression during early imbibition of healthy axes (3 HAI) (Fig. **6c**,**7a,8,9**), further suggesting that axes that failed to germinate have slow or arrested metabolism. Future studies will examine no- and low-GP axes at times shorter and longer than 24 HAI to distinguish these scenarios. Slowed metabolism as a consequence of aging provides an opportunity for “rescue” by extending geminating times and preventing microbial invasion.

### Adding insult to injury

Pinpointing the lethal event is confounded in seeds because damage continues to accumulate postmortem. Most 1996H seeds died by 2013 (Walters *et al.*, 2020), meaning that damage measured in 2018 was beyond the lethal event. The low RIN values in dry seeds of this cohort reflect degraded ribosomal subunits (Schroeder *et al.*, 2006), though some intact ribosomes appear present based on identifiable 18S and 25S peaks in electropherograms (**Fig. S3**). Poor recovery of RNA integrity 24 HAI in 1996H suggests that translation is also impaired, and *de novo* protein synthesis is needed to complete germination (Rajjou *et al.*, 2008). Possibly, imbibition time courses of proteomic changes in healthy vs. aged seeds would illuminate the relationship between translational competence and overall metabolic vigor and radicle emergence.

Lethal events were occurring in the 1999H cohort at the time RNA was extracted in 2018 (Walters et al, 2020). However, damage to rRNA (RIN values < 7) was not obvious in this cohort for dry or imbibed axes (Fig. **2**,**S2,S3**, Table **S10**). In other words, some 1999H axes experienced a “death blow,” but it was not detected by loss of RNA integrity per se (Fleming *et al.*, 2017, 2018). More likely the lethal event is reflected by languishing metabolism upon hydration (Fig. **6b**).

### Are low-GP axes dead or alive?

The low-GP category (Fig. **3**) includes axes from both 2015H and 1999H (98-99% and 60-90% germination, respectively), the latter having an approximate 2:1 ratio of green and white cotyledons after 72 hours of light. Transcriptomes of the readily-identified high-GP group likely clustered separately from the low-GP group because of less intense up- and down-regulation in low-GP axes (Fig. **6**). Low-GP axes shared over 90% of their SDE genes with high-GP axes, which were expressed in the same direction but with higher log_2_FC values (note R^2^ = 0.94 and slope = 1.27 > 1, P << 0.01 in Fig. **5a**). The low-GP category may be more aptly named “*s*low germination potential” to highlight the similarity in expression patterns with healthy axes 6 or 12 HAI (Bellieny-Rabelo *et al.*, 2016) (Fig. **8**). Since time required for radicle emergence varies, from 22-36 HAI for 2015H to 22-50 HAI for 1999H seeds (Fig. **1a**), the low-GP category may encompass the later-germinating seeds from each cohort. Low-GP seeds also may or may not be fully capable of completing germination, as evidenced by the mixture of white and green cotyledons associated with these axes. In low-GP seeds that successfully germinate, delayed transcription may catch up to match high-GP axes if given sufficient imbibition time (Waterworth *et al.*, 2019). In other words, low-GP axes may germinate, or not, depending on experimental conditions, giving high uncertainty to exact germination percentages as well as greater probability of type 1 statistical errors (accepting a seed is dead when it is not (**Fig. S1**).

Detecting an intermediate class of germination potential argues for a transition between ability and inability to germinate that is more continuous than characterizations of alive and dead or green and white cotyledons. Transcriptome size, coordinated gene expression, RNA integrity, and the extent that radicle emergence is delayed may provide usable signals that *quantify* damage in aging seeds before germination capacity is entirely lost. Assessments of seeds that are dying but retain some possibility of surviving are likely to reveal other cellular signals too. Quantitative metrics of damage are needed to phenotype segregating populations of seeds that are aging rapidly or slowly. Observing an intermediate class also provides a new framework to adjust our notions of how dry seeds die, away from catastrophic events and towards slow attrition and eventual death. Here, we find no *single* molecular failure that signals mortality. However, a threshold of metabolic competence may ultimately separate an embryo that cannot be revived from one that can, when given substantial intervention to prolong imbibition and retard microbial growth.

## Conclusion

Dry seeds are neither overtly alive nor dead. Imbibition initiates the process of germination, a complex developmental cascade of fluctuating gene expression that culminates (*sensu stricto*) in radicle emergence. Germination-related transcripts are detected in lethally aged seeds and self-destruct pathways are *not*, ruling out a clear metabolism-based “death signature”. Rather, the distinction between alive and recently dead appears to be kinetic. That is, dead seeds fail to muster a sufficient transcriptome response before microbial infestations dominate. Therefore, global gene expression may serve as an excellent “canary in the coal mine” to indicate seed health or imminent demise.

## Acknowledgements

MBF was supported through the ARS postdoctoral fellowship program. USDA is an equal opportunity employer and provider.

## Author contributions

Design of the research: MBF, ELP, CW; performance of the research: MBF; data analysis and interpretations: MBF, ELP, CW; writing the manuscript: MBF, ELP, CW.

## Data availability

The data from the storage time experiment that support the findings of this study are openly available in the SRA database at ncbi.nlm.nih.gov/sra, reference number PRJNA675850. The data from the imbibition time experiment that support the findings of this study were derived from resources in the public domain, available at the SRA database at ncbi.nlm.nih.gov/sra, reference number PRJNA326110, or in the GEO database at ncbi.nlm.nih.gov/geo/, accession number GSE83481.

## Supporting Information

**Figure S1** Reliability of the cotyledon greening assay in predicting germination capacity of soybean (cv. ‘Williams 82’) axes used for transcriptome sequencing.

**Figure S2** Electropherograms as well as associated cotyledons for total RNA extracted from soybean (cv. ‘Williams 82’) axes from 2015H and 1999H cohorts imbibed for 24 hours.

**Figure S3** Electropherograms for total RNA extracted from soybean (cv. ‘Williams 82’) axes from the 1996H cohort, as well as the cotyledons associated with those axes imbibed for 24 hours.

**Figure S4** Number of transcripts significantly differentially expressed in soybean (cv. ‘BRS 284’) axes imbibed for 3, 6, 12, and 24 hours, when compared to dry axes.

**Table S1** All transcripts from the soybean v.4 transcriptome and their MapMan bin assignments from Mercator4 v.2, used to make Fig. 6,7,9

**Table S2** RIN of each soybean (cv. ‘Williams 82’) embryonic axis used for transcriptome sequencing

**Table S3** Counts for all transcripts in the soybean v.4 transcriptome from soybean (cv. ‘Williams 82’) dry and imbibed axes harvested in 2015, 1999, and 1996 (‘storage time experiment’)

**Table S4** Differential expression of all expressed transcripts in soybean (cv. ‘Williams 82’) imbibed axes from each germination potential category, compared to all dry ‘Williams 82’ axes

**Table S5** Counts for all transcripts in the soybean v.4 transcriptome from soybean (cv. ‘BRS 284’) axes imbibed for 0, 3, 6, 12, and 24 hours (‘imbibition time experiment’)

**Table S6** Differential expression of all expressed transcripts in soybean (cv. ‘BRS 284’) axes from each imbibition time point, compared to all dry ‘BRS 284’ axes

**Table S7** Transcript ID, MapMan annotation, and cluster assignment for all soybean transcripts in each of the six clusters identified in Fig. **8**

**Table S8** Overrepresentation analysis of MapMan annotations found in each of the six clusters identified in Fig. **8**

**Table S9** Annotation, transcript ID, and log_2_FC of all 5760 soybean ‘germination transcripts’ identified from the literature

**Table S10** Normalized log_2_FC values of all soybean ‘germination transcripts’ significantly differentially expressed in at least one germination potential category, used to generate Fig. **9**

