## Supplemental Information for "Too little, too late: transcription during imbibition of lethally aged soybean seeds is weak and delayed, but not aberrant"

### *New Phytologist* Supporting Information

Article acceptance date: Click here to enter a date.

The following Supporting Information is available for this article:

**Fig. S1** The reliability of the cotyledon greening assay in predicting germination capacity of soybean (cv. ‘Williams 82’) axes used for transcriptome sequencing: Page 3

**Fig. S2** Electropherograms as well as associated cotyledons for sequenced soybean (cv. ‘Williams 82’) axes from 2015H and 1999H cohorts imbibed for 24 hours: Page 4

**Fig. S3** Electropherograms for all sequenced soybean (cv. ‘Williams 82’) axes from the 1996H cohort, as well as the cotyledons associated with those axes imbibed for 24 hours: Page 5

**Fig. S4** Number of transcripts significantly differentially expressed in soybean ‘BRS 284’ axes imbibed for 3, 6, 12, and 24 hours, when compared to dry axes: Page 6

**Table S1** Counts for all transcripts in the soybean v.4 transcriptome from soybean (cv. ‘Williams 82’) dry and imbibed axes harvested in 2015, 1999, and 1996 (‘storage time experiment’): Given in a separate excel file

**Table S2** Counts for all transcripts in the soybean v.4 transcriptome from soybean (cv. ‘BRS 284’) axes imbibed for 0, 3, 6, 12, and 24 hours (‘imbibition time experiment’): Given in a separate excel file

**Table S3** Differential expression of all expressed transcripts in soybean (cv. ‘Williams 82’) imbibed axes from each germination potential category, compared to all dry ‘Williams 82’ axes: Given in a separate excel file

**Table S4** Differential expression of all expressed transcripts in soybean (cv. ‘BRS 284’) axes from each imbibition time point, compared to all dry ‘BRS 284’ axes: Given in a separate excel file

**Table S5** All transcripts from the soybean v.4 transcriptome and their MapMan bin assignments from Mercator4 v.2, used to make Fig. 6,7,9: Given in a separate excel file

**Table S6** Transcript ID, MapMan annotation, and cluster assignment for all soybean transcripts in each of the six clusters identified in Fig. **8**: Given in a separate excel file

**Table S7** Overrepresentation analysis of MapMan annotations found in each of the six clusters identified in Fig. **8**: Given in a separate excel file

**Table S8** Annotation, transcript ID, and log_2_FC of all 5712 soybean ‘germination transcripts’ identified from the literature: Given in a separate excel file

**Table S9** Normalized log_2_FC values of all soybean ‘germination transcripts’ significantly differentially expressed in at least one germination potential category, used to generate Fig. **9**: Given in a separate excel file

**Table S10** RIN of each soybean (cv. ‘Williams 82’) embryonic axis used for transcriptome sequencing: Page 7

**Fig. S1** Correspondence of cotyledon greening and excised embryonic axis expansion in soybean seeds from three cohorts. Both greening and expansion assays gave comparable percentages as standard germination assays, and allowed us to predict viability of axes used for transcriptome sequencing from the greening response of the associated cotyledon. Light grey quadrants depict incidence of type I and type II errors. We did not see a seed that had greening cotyledons and an inviable axis; about 8% of 1999H axes expanded even though the corresponding cotyledon did not green in the allotted 3 day period.

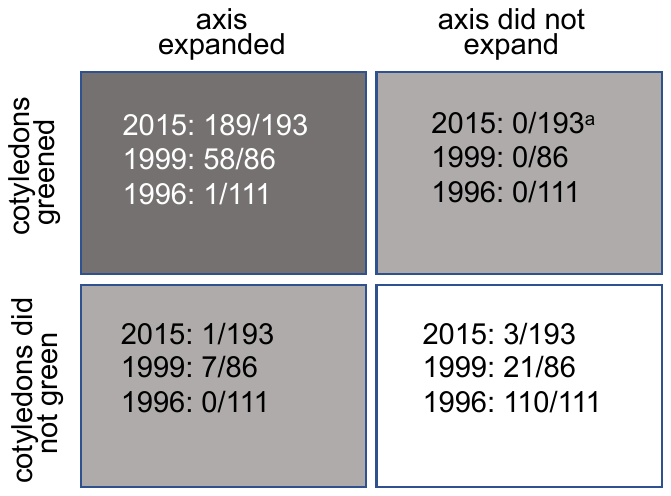

**Fig. S2** Electropherograms as well as associated cotyledons for total RNA extracted from soybean (cv. ‘Williams 82’) axes from 2015H and 1999H cohorts imbibed for 24 hours. Top row: 1999H axes for which corresponding cotyledons did not green (see photograph). Middle row: electropherograms from 1999H axes for which corresponding cotyledons greened. Bottom row: electropherograms of 2015H axes in which 98% germinated and greened; red arrow indicates plumule expansion. The 5 panels boxed in red are from axes that clustered into the low-GP group in Fig. **3**.

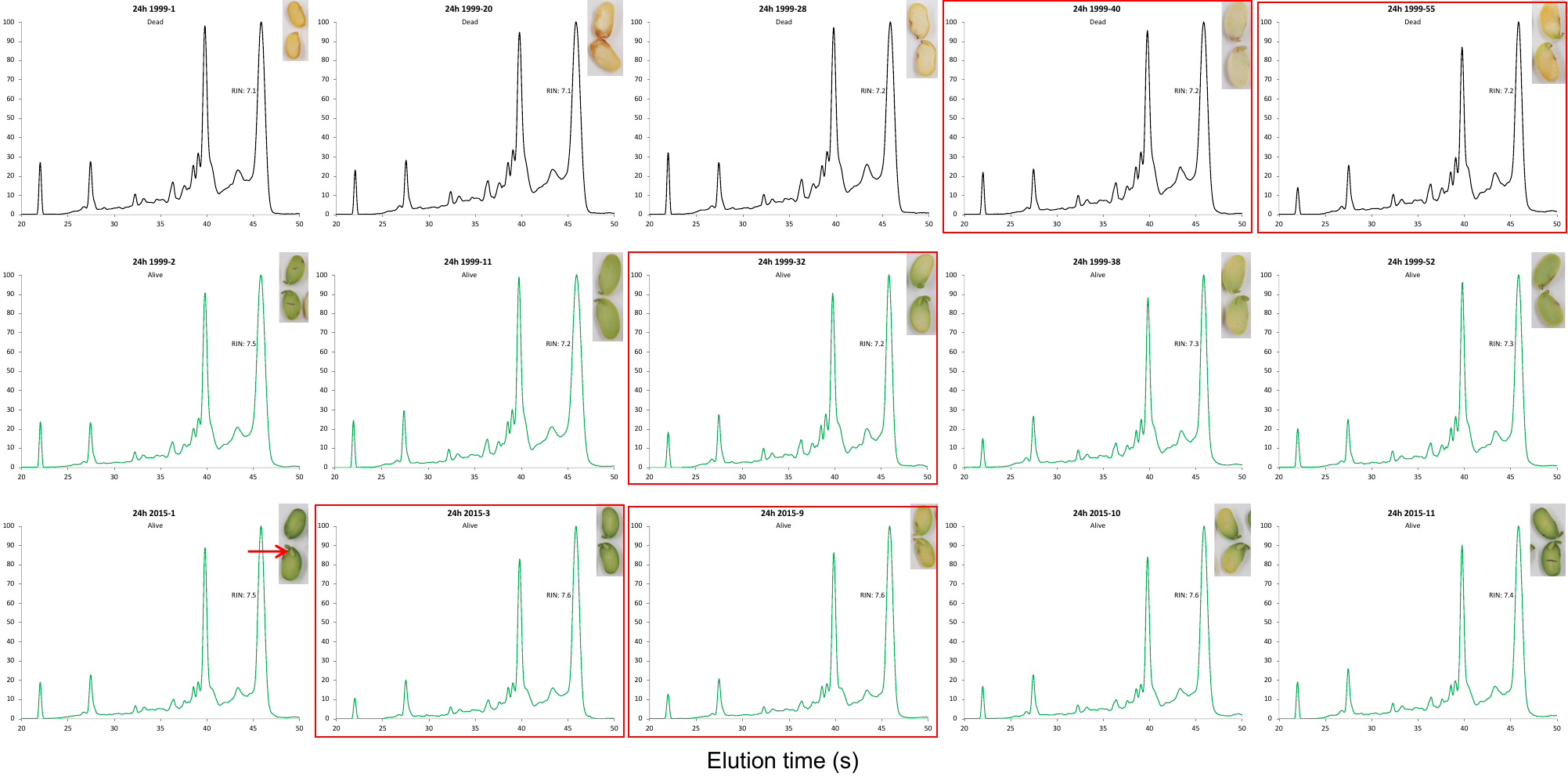

**Fig. S3** Electropherograms for total RNA extracted from soybean (cv. ‘Williams 82’) axes from the 1996H cohort, as well as the cotyledons associated with those axes imbibed for 24 hours. RNA was extracted from 1996H axes that were dry (top row) and imbibed 24 hours (bottom row). Cotyledons from imbibed seeds did not green and clustered in the no-GP group, consistent with a measured 1% germination (Fig. **1a**). RIN values for these electropherograms averaged 5.7.

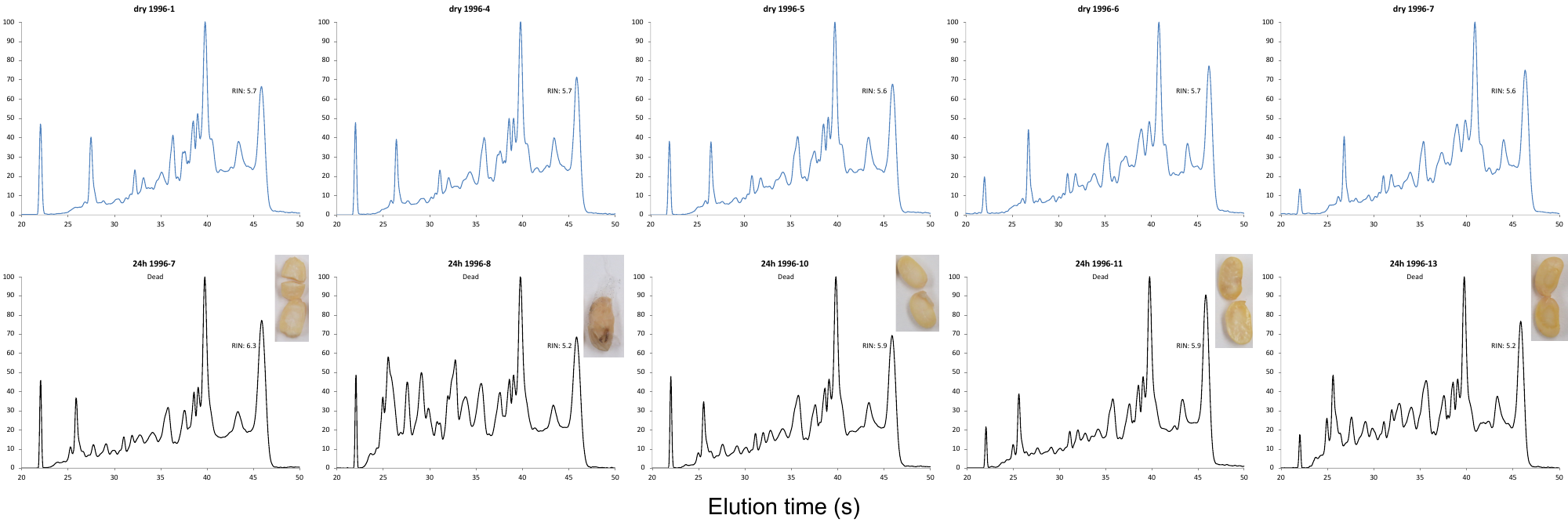

**Fig. S4** Venn diagram of transcripts significantly (|log_2_FC| > 2 and p < 0.001) differentially expressed in axes from soybean (cv. ‘BRS 284’) at different hours after imbibition (HAI) compared to dry axes. A total of 29,701 genes were differentially expressed in at least one imbibition time-point. Axes 24 HAI had the largest number of differentially expressed transcripts, and axes 3 HAI had the fewest. The majority of transcripts differentially expressed in axes 3 HAI were identified in all imbibition time-points. Axes 6 and 12 HAI had an intermediate number of differentially expressed transcripts, most of which were shared with the other time-points.

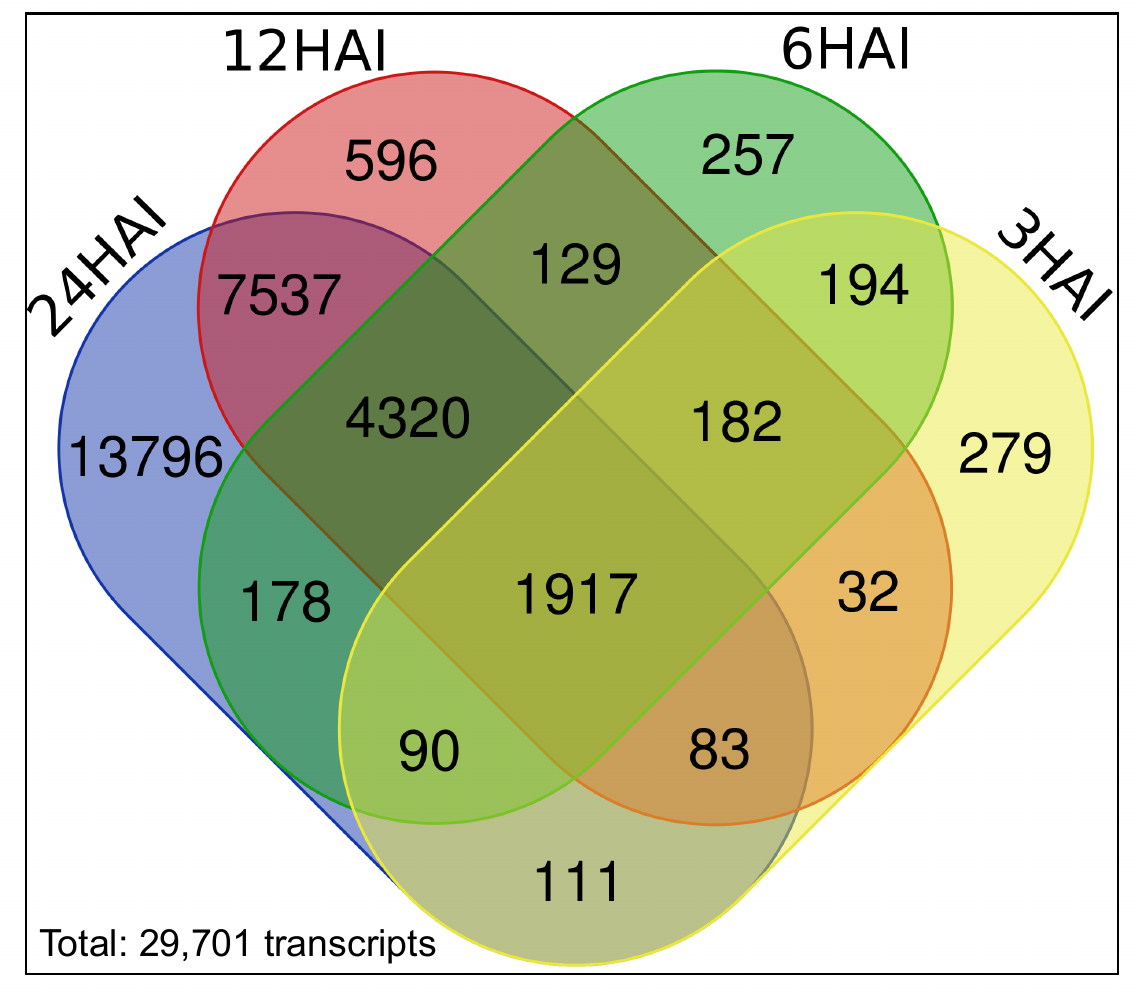

**Table S10** RIN of each soybean embryonic axis used for transcriptome sequencing. Axes were removed from either dry seeds or seeds after 24 hours of imbibition. Cotyledons of imbibed seeds were placed in the light and observed for color changes after three days. Mean RINs with the same connecting letter are not significantly different from each other at alpha = 0.05 (Tukey’s HSD). SD = standard deviation.

| Harvest year | Cotyledon color | RIN | | | | | mean RIN ± SD | Connecting |
| --- | --- | --- | --- | --- | --- | --- | --- | --- |
|  |  | axis 1 | axis 2 | axis 3 | axis 4 | axis 5 |  | letters |
| 1996 | dry | 5.7 | 5.7 | 5.6 | 5.7 | 5.6 | 5.66 ± 0.05 | a |
| 1996 | white | 6.3 | 5.2 | 5.9 | 5.9 | 5.2 | 5.70 ± 0.48 | a |
| 1999 | dry | 7.1 | 7.0 | 7.2 | 7.1 | 7.0 | 7.08 ± 0.08 | b |
| 1999 | white | 7.1 | 7.1 | 7.2 | 7.2 | 7.2 | 7.16 ± 0.05 | bc |
| 1999 | green | 7.5 | 7.2 | 7.2 | 7.3 | 7.3 | 7.30 ± 0.12 | bc |
| 2015 | dry | 7.3 | 7.6 | 7.5 | 7.5 | 7.5 | 7.48 ± 0.11 | bc |
| 2015 | green | 7.5 | 7.6 | 7.6 | 7.6 | 7.4 | 7.54 ± 0.09 | c |
